# H- and m-channel overexpression promotes seizure-like events by impairing the ability of inhibitory neurons to process correlated inputs

**DOI:** 10.1101/2024.11.25.625196

**Authors:** Scott Rich, Taufik A. Valiante, Jeremie Lefebvre

**Author notes:** **For correspondence:** (SR).

## Abstract

Channelopathies affecting h- and m-channels present a paradox in epilepsy research: while both over- and underexpression of these channels can be epileptogenic, channel overexpression does not appear to increase the excitatory-inhibitory (E-I) balance as caused by channel underexpression. We here derive a viable mechanism for ictogenesis driven by h- and m-channel overexpression from analysis of an *in silico* spiking neuronal microcircuit exhibiting spontaneous seizure-like events (SLEs). Such SLEs are dependent upon sufficiently strong gain in two adaptation terms phenomenologically modeling these channels’ effects: voltage homeostasis (h-current) and spike-frequency adaptation (m-current). Excessive gain of these adaptation terms interferes with the circuit’s processing of highly correlated input, promoting a sequence of network-level events that collectively provoke an SLE. Importantly, these changes do not cause increased excitability in isolated neurons, nor does this cascade require a change in the amplitude of external input to the circuit, suggesting an ictogenic pathway independent of classical changes to the E-I balance. The viability of this mechanism for SLE onset is strengthened by the host of experimentally-characterized features of seizure produced in this model reliant upon the presence of these adaptation terms, including the irregular initiation and termination of seizure-like events and time-varying peak frequency of oscillations during such events (i.e., chirps). Moreover, the cell-type dependent effects of changes in these adaptation terms, as delineated in our analyses, represent experimentally-testable predictions for future study of h- and m-channelopathies. These computational results provide vital new insights into the epileptogenic nature of h- and m-channel overexpression currently absent in the experimental literature.

## Introduction

Epilepsy, the most common serious neurological disorder in the world (***Reynolds, 2002***), is typified by the brain’s predisposition to repeated and unpredictable transitions into hypersynchronous and hyperactive states called seizures (***Jiruska et al., 2013***). Despite important advances, understanding the myriad etiologies underlying epileptogenesis and ictogenesis remains a formidable challenge (***Jasper, 2012; Dehghani et al., 2016; Žiburkus et al., 2013; Rich et al., 2022; Sutula and Dudek, 2007; Cossart et al., 2001; Cobos et al., 2005; Arnold et al., 2019; Klaassen et al., 2006; Albertson et al., 2011; Rich et al., 2020a; Chang et al., 2018***). Many of these are jointly classified as channelopathies, with the pathological impetus promoting seizure lying in mutations to genes encoding ion channels (***Ng et al., 2024; Mulley et al., 2003; Lascano et al., 2016; Menezes et al., 2020***). Two ion channels of particular interest are the hyperpolarization-activated cation (h-) and m-channels, which drive key homeostatic and adaptive processes in neurons (***Benda and Herz, 2003; Gasselin et al., 2015; Rathour and Kaphzan, 2023; Gasselin et al., 2017; Zhou et al., 2018; Otto et al., 2006***) and have thus emerged as key regulators of neuronal excitability and function. H- and m-channelopathies have been implicated in epileptogenesis (***Lerche et al., 2001; Rogawski, 2000; Steinlein and Noebels, 2000; Byers et al., 2021; Poolos, 2004; DiFrancesco et al., 2019; Noam et al., 2011; Bender et al., 2003; Dyhrfjeld-Johnsen et al., 2009; Arnold et al., 2019***) and control of these channels’ activity has shown promise in seizure prevention (***Chen et al., 2002; Rogawski, 2000; Lerche et al., 2001***).

However, the relationship between these channels and seizure appears surprisingly complex. For example, both over- (***Poolos, 2004; DiFrancesco et al., 2019; Noam et al., 2011; Bender et al., 2003; Dyhrfjeld-Johnsen et al., 2009***) and under- (***Poolos, 2004; DiFrancesco et al., 2019; Arnold et al., 2019; Dyhrfjeld-Johnsen et al., 2009***) expression of the h-channel can be epileptogenic. While the effects of h-channel underexpression are intuitively understood through effects on a neuronal population’s excitatory-inhibitory (E-I) balance, mechanisms explaining the epileptogenic effects of h-channel overexpression are relatively understudied. Similarly, while decreased expression of the m-channel is more commonly associated with epilepsy (***Lerche et al., 2001; Rogawski, 2000; Steinlein and Noebels, 2000; Byers et al., 2021***) given the m-current’s role in driving spike-frequency adaptation (***Peters et al., 2005; Benda and Herz, 2003; Gu et al., 2005***), recent studies have identified epileptogenic gain of function mutations related to the m-channel (***Niday and Tzingounis, 2018***). Mechanistic explanations for the less intuitive epileptogenic effects of h- and m-channel overexpression could yield new therapeutic targets for the approximately one-third of epilepsy patients who do not respond to current pharmaceutical interventions (***Brodie and Sills, 2011; Kwan et al., 2001***).

*In silico* tools are ideally suited to decipher these relationships, as a host of computational studies (***Wilson and Cowan, 1972; Kramer et al., 2005; Yang et al., 2005; Lytton, 2008; Cressman et al., 2009; Ullah et al., 2009; Kramer et al., 2012; Jirsa et al., 2014; Wendling et al., 2016; Buchin et al., 2016; Chizhov et al., 2018; Rich et al., 2020a; El Houssaini et al., 2020; Liou et al., 2020; Rich et al., 2020b, 2022; Depannemaecker et al., 2021***) have described detailed dynamical and physiological mechanisms underlying epileptiform activity. Here, we build on these approaches and our previous work (***Rich et al., 2020b, 2022***) to explore the role of h- and m-channel misexpression in seizure susceptibility. Specifically, we built a spiking E-I neuronal microcircuit accounting for activity driven by the h- and m-channels by endowing each neuron with a phenomenological voltage homeostasis current (reflecting the impact of h-current on neuronal activity (***Gasselin et al., 2015***) and therefore referred to as h-adaptation) and a spike-frequency adaptation term (echoing a primary role of the m-current (***Benda and Herz, 2003; Gu et al., 2005; Peters et al., 2005***) and therefore referred to as m-adaptation). Our previous work showed that this network architecture (without adaptation) would suddenly transition into hypersynchronous and hyperactive dynamics via a saddle-node bifurcation (***Chow and Hale, 2012; Rich et al., 2021, 2020b***) with increasing amplitude of a tonic external input. Here, we expose the new model to dynamic stimuli in the form of noisy inputs with varying correlation, emulating fluctuations arising naturally due to recurrent circuit connectivity (***Kriener et al., 2008***), synchrony (***Roelfsema et al., 2004***) and/or common afferent inputs including sensory stimuli (***Middleton et al., 2012; Butler et al., 2017; Doiron et al., 2016; Honey and Valiante, 2017; Lee et al., 1998; Stark et al., 2008***). Such inputs interrogate the network’s stability and, in turn, susceptibility to seizure-like events (SLEs).

Our analyses show that sufficiently strong h- and m-adaptation impairs the ability of this network to process correlated inputs, which manifests in more frequent spontaneous (***Kuhlmann et al., 2018; Mormann et al., 2007***) transitions into and out of SLEs, outlining a key role for excessive h- and m-adaptation in seizure onset and termination. Furthermore, these adaptation terms were found to drive non-stationary oscillatory behavior commonly observed during seizure (i.e., chirps or “glissandi” (***Schevon et al., 2012; Cunningham et al., 2012***)), in which neural activity traverses multiple dynamical regimes characterized by distinct spectral properties (***Avoli and de Curtis, 2011***). We precisely discerned how increased gain in the h- and m-adaptation affects network activity and increases the system’s vulnerability to SLEs: the unique sensitivity to input correlation of systems with sufficiently strong h- and m-adaptation promotes high amplitude fluctuations in inhibitory activity. SLEs arise when the inhibition resulting from such fluctuations is translated into post-inhibitory hyperexcitability in the excitatory population via excessive h-adaptation, combined with the inability of inhibitory neurons to restrain this hyperexcitability due to excessive m-adaptation. This mechanism additionally provide support for the “paradoxical” effects of h- and m-channelopathies on seizure susceptibility via these epileptogenic changes’ dependence on cell type: the epileptogenic effects of gain of function mutations promoting m-channel expression (***Niday and Tzingounis, 2018***) can be explained by increased spike-frequency adaptation in inhibitory neurons overshadowing similar effects in excitatory neurons, and similarly the epileptogenic effects of h-channel overexpression (***Poolos, 2004; DiFrancesco et al., 2019; Noam et al., 2011; Bender et al., 2003; Dyhrfjeld-Johnsen et al., 2009***) may arise when post-inhibitory excitability in excitatory cells overshadows analagous changes to inhibitory neurons. These results provide important mechanistic insights into the epileptogenic effects of h- and m-channelopathies via a new *in silico* seizure model reflecting the diverse etiologies of epilepsy.

## Results

### Excessive voltage homeostasis and spike-frequency adaptation promotes realistic *in silico* seizure-like events

In the computational study of epilepsy, particularly when using spiking neuronal microcircuits (***Rich et al., 2020b, 2022; Liou et al., 2020***), *in silico* approximations of seizure commonly focus on capturing transitions in and/or out of hypersynchronous and hyperactive neuronal activity states (***Jiruska et al., 2013; Kramer et al., 2005, 2012***). However, additional salient dynamical features of seizure dynamics differentiate these oscillations from ones serving physiological functions. One key differentiator is the non-stationary spectral characteristics of ictal activity that change over time in a stereotyped manner (***Avoli et al., 2016; Florez et al., 2013***). This pattern, characterized by a ramping up and slowing down of the peak frequency of ictal activity, is oftentimes termed chirps or “glissandi” (***Schevon et al., 2012; Cunningham et al., 2012***). Additionally, the most commonly classified seizure subtype, low-voltage fast-onset (LVF) seizures (***Lee et al., 2000; Avoli et al., 2016; Perucca et al., 2014***), is noted for the seeming complicity of inhibitory neurons in seizure onset (***Shiri et al., 2015; Elahian et al., 2018; Weiss et al., 2019***) and initial peak oscillatory frequencies in the 10-30 Hz range (***Florez et al., 2013; Jiruska et al., 2013; Elahian et al., 2018; Velascol et al., 1999; Lee et al., 2000; Weiss et al., 2019; Perucca et al., 2014; Wendling et al., 2005***).

Building on an E-I network architecture (diagram in Figure 1**A**) widely used to model both oscillatory and epileptiform activity (***Wilson and Cowan, 1972; Kramer et al., 2005; Lytton, 2008; Kramer et al., 2012; Wendling et al., 2016; Rich et al., 2020b, 2022; Depannemaecker et al., 2021***), we developed a model in which the addition of h- and m-adaptation promotes spontaneous, irregular and unpredictable (***Kuhlmann et al., 2018; Mormann et al., 2007***) transitions into SLEs exhibiting these distinguishing fluctuating spectral features of seizure (see details in the Materials and Methods). Figure 1**A** showcases one exemplar SLE generated by this model, which includes pronounced voltage homeostasis and spike-frequency adaptation via high gains of the h- and m-adaptation terms; the full simulation yielding this SLE, decomposed into its excitatory and inhibitory activity, is shown in Figure 1**B**. For a majority of the 200 second simulation, activity remains asynchronous (a state we will later refer to as the model’s “baseline”), although brief instances of synchrony reminiscent of inter-ictal spikes (***Staley and Dudek, 2006***) are observed. Two SLEs are observed at approximately 40 and 70 seconds, both of which terminate within 10 seconds. The frequency profiles of excitatory activity in these hyperactive states (second row, Figure 1**B**) exhibit a peak frequency just below 20 Hz and dynamics reminiscent of chirps. This is more clearly visualized by zooming in on the second SLE in Figure 1**C**. Our confidence in the classification of these events as SLEs is strengthened by the correspondence between these spectrograms and analogous experimental recordings of ictal events in human cortical (***Florez et al., 2013***) and subicular (***Huberfeld et al., 2015***) tissue, intracerebral EEG from human hippocampus (***Wendling et al., 2005***), and EEG from non-human primates (***Schellenberg and Devergnas, 2020; Vuong et al., 2020***). This alignment with experimental data was further confirmed via quantification of the intervals between SLEs (Supplementary Figure S1), which are well fit by a gamma distribution—this matches experimental quantifications of interictal distributions that also tend to be gamma distributed (***Breton et al., 2020; Shayegh et al., 2014; Suffczynski et al., 2006; Snyder et al., 2008; Bauer et al., 2017; Colic et al., 2013***).

**Figure 1.**
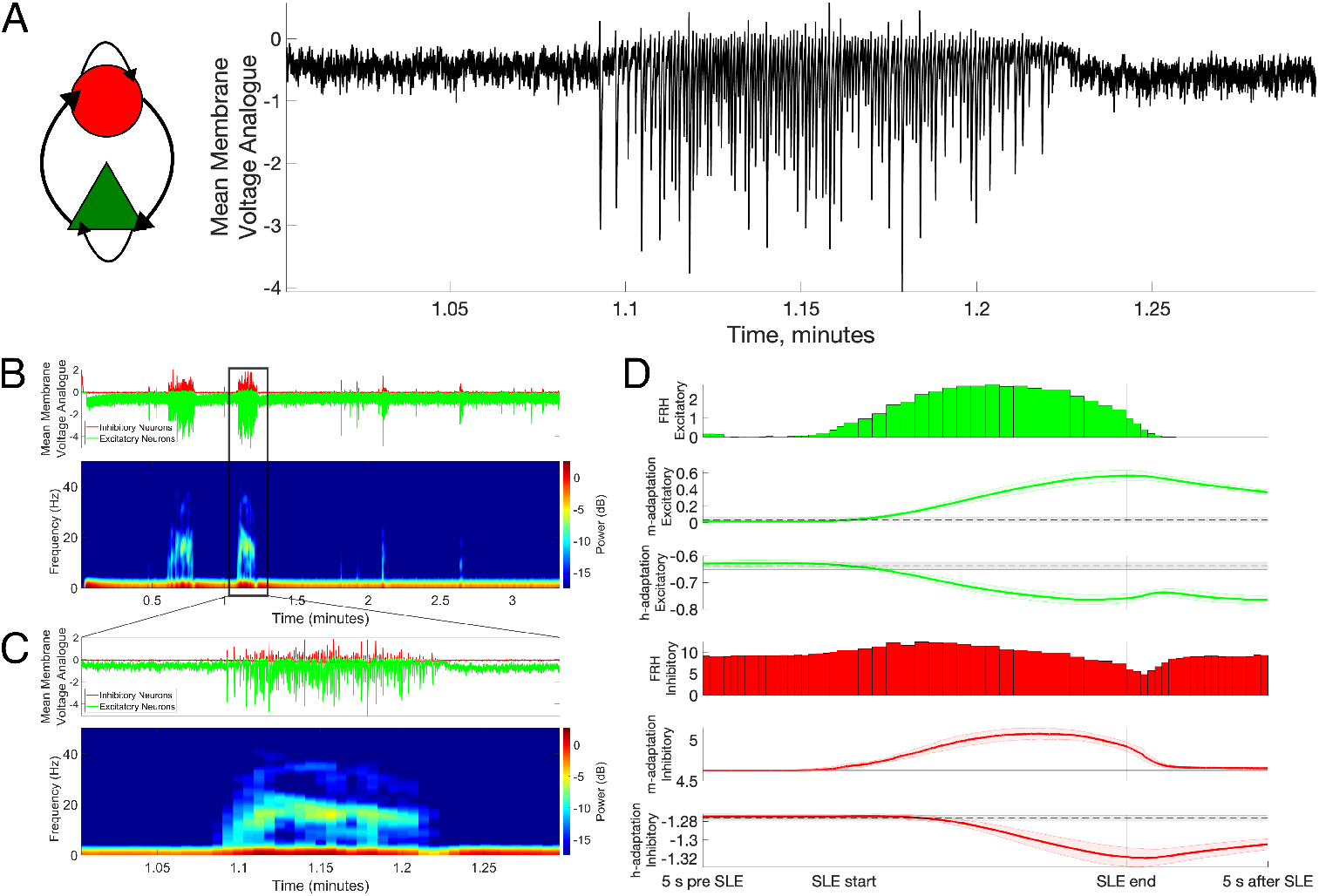
E-I microcircuit reproduces key characteristics of seizure dynamics driven by neuronal adaptation. **A**: The E-I network presented here (diagrammed at left, see Materials and Methods for details) spontaneously and stochastically transitions between periods of sparse, asynchronous activity and seizure-like periods of high-frequency activity when subjected to an input with correlation *c* = .10 (an example is shown at right). Mean membrane voltage analogue is the mean value of the variable *u* taken over all excitatory and inhibitory neurons and is a rough analogue of an experimental local field potential (LFP). **B**: Dynamics of the full simulation yielding Panel **A** separated between excitatory and inhibitory populations (first row) along with a spectrogram of mean activity of the excitatory population (second row). **C**: A more precise view of a single seizure-like event. Panels are as in **B**. Apparent in this visualization is a peak frequency of the seizure-like dynamics in the 10-30 Hz range and the chirp (ramp up and ramp down) of peak frequency during the seizure-like event. **D**: Average spiking dynamics and activity of the voltage homeostasis current (h-adaptation) and spike-frequency adaptation term (m-adaptation) around 57 spontaneously arising SLEs over 100 independent simulations, separated between excitatory and inhibitory neurons. Spiking dynamics quantified via discrete firing rate histograms (FRHs), while the population averaged activity of the h- and m-adaptation are averaged over the normalized duration of each SLE (faint shading represents ± one SD). Black dotted line (with faint shading representing ± one SD) represents the mean of these terms outside of SLEs. Apparent in this visualization is the increase in inhibitory activity above the mean prior to SLE onset (via increased inhibitory m-adaptation prior to SLE start), the resulting increase in the magnitude of excitatory h-adaptation around SLE start, and the termination of the SLE associated with maximal excitatory m-adaptation (peak in excitatory m-adaptation around SLE end).

To delineate the role of voltage homeostasis and spike frequency adaptation in these events, we captured the trajectory of the h- and m-adaptation terms during SLEs in both the excitatory and inhibitory populations. We normalized and averaged h- and m-adaptation time series around 57 SLEs to yield a visualization of these terms’ typical progression before, during, and after an average SLE in Figure 1**D** (see details in Materials and Methods). Immediately apparent in these plots are important changes in h- and m-adaptation from the moments immediately preceding an SLE through its termination, in contrast with their near constant values in the 5 seconds prior to SLEs. Notable in these trajectories are the following key features: 1) As evidenced by increased m-adaptation (which tracks spiking activity with a time delay; see Materials and Methods for further details), the first deviation from mean non-SLE activity is increased inhibitory cell activity preceeding SLE onset; 2) This increased inhibitory activity hyperpolarizes the excitatory population; 3) The excitatory h- adaptation (which tracks voltage with a time delay; see Materials and Methods for further details) builds up during this hyperpolarization; 4) This translates into a net increase in the excitability of excitatory neurons in the moments around SLE onset; 5) Inhibitory cells are unable to compensate for the increase in excitatory excitability hallmarking the SLE due to rapid increases in the m-adaptation; and 6) SLE end corresponds with a peak in the excitatory m-adaptation, indicating that excitatory spike-frequency adaptation is responsible for SLE termination.

This analysis showcases a potential complicit role of h- and m-adaptation, with cell-type specific effects, in the onset of SLEs. Given this, we would also expect these terms to have some influence on network dynamics more generally, which we assessed via network activity as a function of the gains dictating the strength of each adaptation term independently in the excitatory 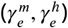 and inhibitory 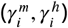 populations. These analyses, presented in Figure 2, reveal the relative importance of inhibitory m-adaptation 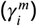 and excitatory h-adaptation 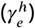 in modulating network activity, both during SLE and non-SLE periods. Indeed, increases to 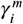 (Figure 2**B**) cause notable suppression of the inhibitory firing rate both inside and outside of SLEs, and corresponding amplification of the excitatory firing rate. Similarly, increases in 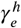 (Figure 2**C**) contribute to notable increases in excitatory activity both inside and outside of SLEs, although with limited effect on inhibitory firing.

**Figure 2.**
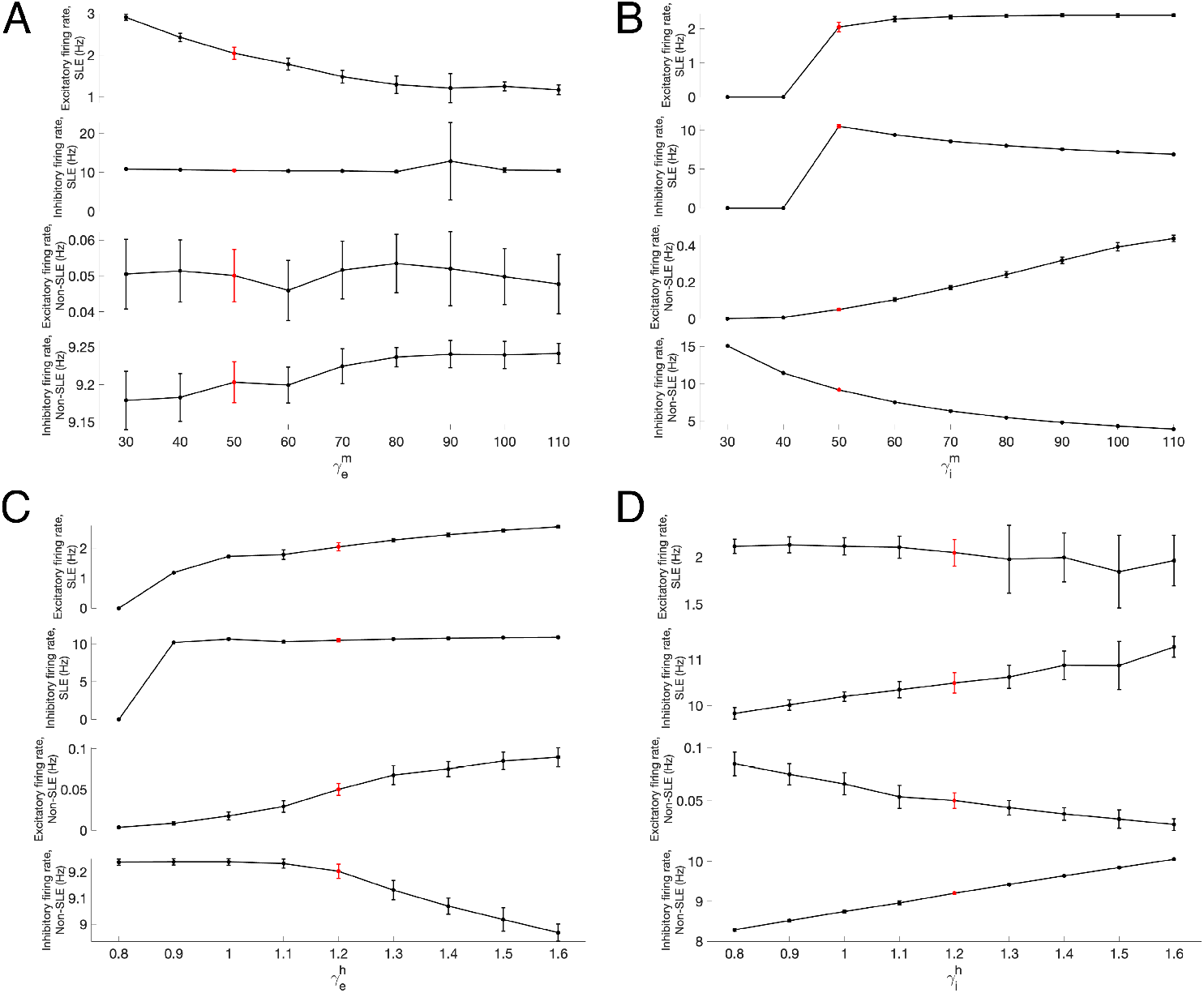
Effects of h- and m-adaptation on firing rates and excitability of excitatory and inhibitory cells inside and outside of SLEs. **A-D**: Modulation of excitatory firing rate during SLEs (first row), inhibitory firing rate during SLEs (second row), excitatory firing rate outside of SLEs (third row), and inhibitory firing rate outside of SLEs (bottom row) as the four adaptation terms 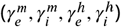 are varied independently. The red point represents the default model parameters (note that the varied y-axes to explain the different error bars in each panel). Firing rates of 0 during SLEs (in Panels **B** and **C**) are indicative of no SLEs occurring for that parameter value. Plots show mean ± one SD over 25 independent 200 s simulations. Changes in 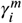 (Panel **B**) and 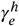 (Panel **C**) have the most pronounced effect on network dynamics, particularly via a monotonic increase in non-SLE excitatory firing rate as these weights are increased. The most pronounced effect of 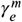 (Panel **A**) is a decrease in excitatory firing rate during SLEs, while the most pronounced effect of 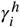 (Panel **D**) is increased inhibitory firing rates throughout the simulation.

In contrast, changes to 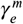 (Figure 2**A**) and 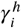 (Figure 2**D**) have little to no effect on firing rates of either excitatory or inhibitory cells. These results showcase that increased 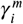 and 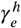 may indirectly increase collective neuronal activity—interfering with E-I balance in this ictogenic microcircuit—in fashions driven by network effects rather than their influence on single-cell excitability.

### Cell-type specific overexpression of h- and m-adaptation increases SLE susceptibility

The activity of the h- and m-adaptation before and during SLEs is evidence that these sources of adaptation not only are necessary to capture key nuances of seizure dynamics, but potentially serve a role in the initiation of these events. This hypothesis is further supported by the results of Figure 2, particularly the increased network activity seen as a function of 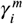 and 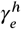 We validated this hypothesis by explicitly quantifying the SLE rate (SLEs per second as calculated over a 200 s simulation) as a function of these gain terms, noting the higher rates than one would expect in a patient with epilepsy or an animal model of the disorder are reflective of a conscientious choice to create a model hyper-vulnerable to SLEs (facilitating efficient *in silico* experiments given computational limitations). We first varied the gain of the h- and m-adaptations uniformly for the excitatory and inhibitory populations (Figure 3**A**), revealing that SLE rate is non-linearly dependent on h- and m-adaptation gain: the effect of 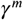 on SLE rate is non-monotonic, while the effect of 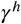 varies largely depending on 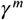 These results motivate the need to disambiguate the effects of these cell-type specific adaptation terms in each neuron population independently.

**Figure 3.**
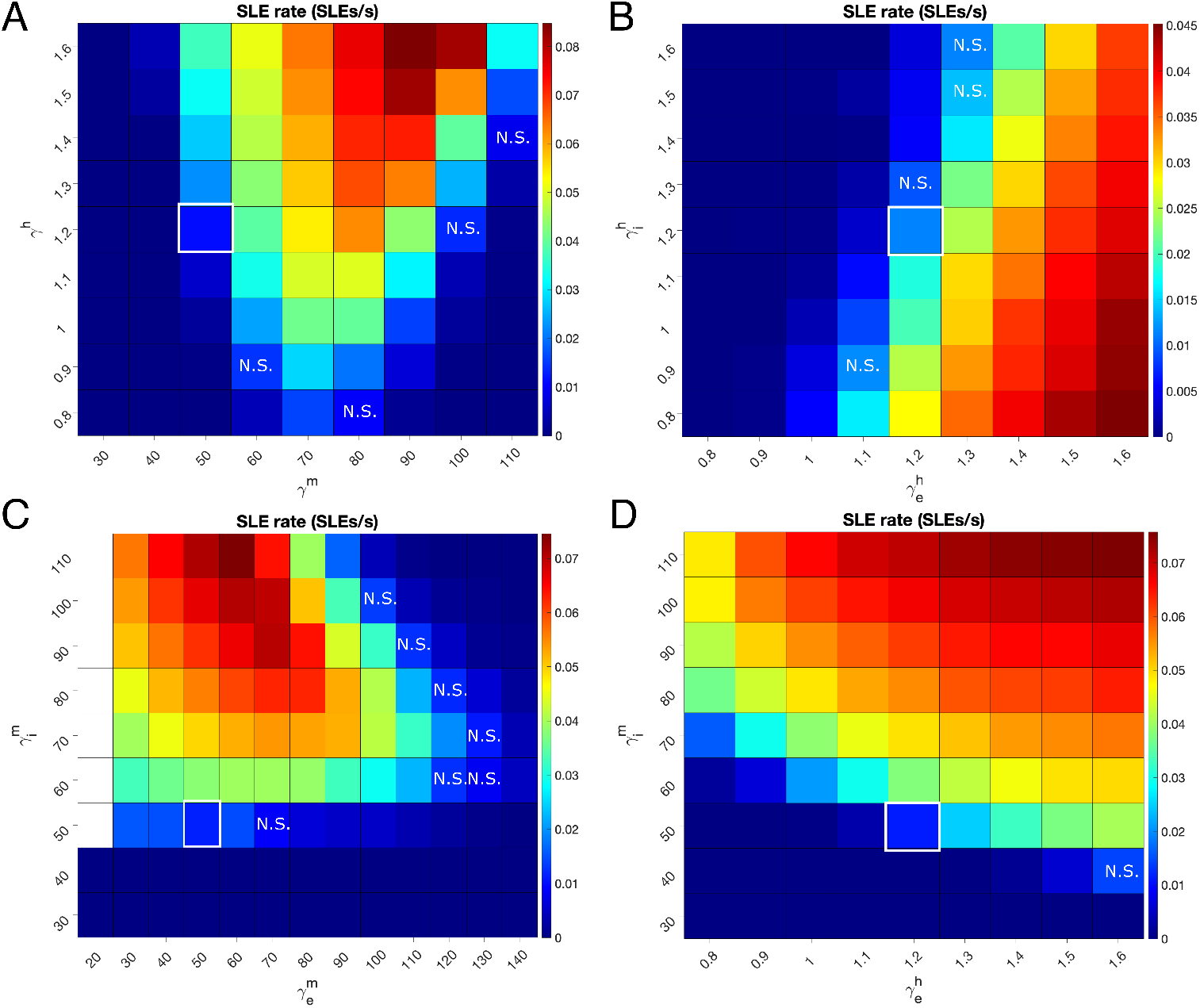
Effects of h- and m-adaptation gains and cell types on seizure-like event (SLE) rates. **A**: Heatmap of SLE rate (SLEs/s) as a function of the h-adaptation gain (γ^*h*^) and m-adaptation gain (γ^*m*^) varied uniformly for the excitatory and inhibitory populations. SLE rate is calculated over a 200 second simulation and averaged over 25 independent trials, with a high input correlation of *c* = .99 to maximize the system’s vulnerability to SLEs (see Figure 4**A**). Default values (γ^*h*^ = 1.2, γ^*m*^ = 50; demarcated by white border) result in an average of approximately two SLEs per 200 second simulation. Increasing *γ*^*m*^ increases SLE rate, while changes to γ^*m*^ have a more complex effect. With few exceptions (squares marked N.S. for non-significant), changes in γ^*m*^ and γ^*h*^ exhibit a significantly different SLE rate from the default values (two-sample t-test,*p* < 0.05). **B**: Heatmap of SLE rate as a function of the h-adaptation gain in the inhibitory 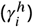 and excitatory 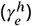 populations. Results as in Panel **A**. Increasing 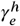 increases SLE rate, while increasing 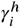 decreases it. Default values demarcated by white border. With few exceptions (squares marked N.S. for non-significant), changes in 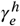 and 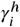 exhibit a significantly different SLE rate from the default values (two-sample t-test, *p* < 0.05). **C**:Heatmap of SLE rate as a function of the m-adaptation gain in the inhibitory 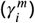 and excitatory 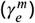 populations. Results as in Panel **A**. Increasing 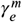 decreases SLE rate, while increasing 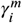 increases it. White squares indicate anomalous networks which do not exhibit termination of seizure-like activity. Default values demarcated by white border. With few exceptions (squares marked N.S. for non-significant), changes in 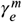 and 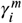 exhibit a significantly different SLE rate from the default values (two-sample t-test, *p* < 0.05). **D**: Heatmap of SLE rate as a function of the m-adaptation gain in the inhibitory population 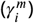 and the h-adaptation gain in the excitatory population 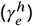 Results as in Panel **A**. Increases in both gains monotonically increase the SLE rate throughout the entire parameter space, supporting the hypothesized key role for these terms in SLE onset. With one exception (square marked N.S. for non-significant), changes in 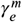 and 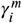 exhibit a significantly different SLE rate from the default values (two-sample t-test, *p* < 0.05).

The gain of the h-adaptation in the excitatory and inhibitory populations was hence independently varied in Figure 3**B**. These analyses reveal that excitatory and inhibitory h-adaptation gains have an opposite impact on SLE rate: while increasing 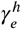 leads to an increased SLE rate, increasing 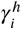 has the opposite effect. This follows from the intuition derived from our analyses of Figures 1 and 2: increased 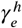 enhances the post-inhibitory hyperexcitability of the excitatory population as a result of hyperpolarization resulting from a period of enhanced inhibitory activity. The monotonic increase in SLE rate as a function of 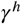 in Figure 3**A**, despite the dichotomous effects of 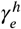 and 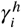 in Figure 3**B**, indicates that the seizure-promoting effect of the h-adaptation in excitatory cells outweighs a reverse effect in inhibitory cells, and further conforms with the analyses of Figures 1 and 2.

Similarly, the effects of the m-adaptation gain are cell-type dependent. Figure 3C shows that increases to 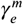 generally reduce the SLE rate (albeit non-monotonically), while the SLE rate increases monotonically with increasing 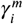 This analysis explains the non-monotonic effect of varying *γ*^*m*^ seen in Figure 3**A**: at low *γ*^*m*^ increased inhibitory activity suppresses SLEs, while at high *γ*^*m*^ decreased excitatory activity suppresses SLEs despite an analogous decrease in inhibitory activity. These results also conform with analysis of Figure 2 highlighting the relative importance of 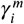 over 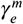 in network activity and SLE susceptibility.

A uniform pattern emerges when we jointly vary the strength of the adaptation terms hypothesized to most directly affect SLE rate: 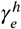 and 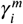 Figure 3**D** shows that increases in both these terms monotonically increase the SLE rate, conforming with the hypothesized culpability of excitatory h-adaptation and inhibitory m-adaptation in SLE onset derived from analyzing Figures 1 and 2. Collectively, these results reveal that the ictogenic potential of h and m-adaptation overexpression is directly linked to cell type, indicating that both over- and under-expression of these channels can be ictogenic via two distinct mechanisms promoting seizure onset.

### H- and m-channelopathies promote SLE susceptibility by impairing the processing of correlated input

With the significant, cell type-specific effect of h- and m-adaptation gain on SLE rate established, we next sought a mechanistic explanation for this phenomenon that could serve to reconcile the paradoxical relationship between h- and m-channelopathies and epilepsy. To do so, we interrogated the model using a correlated input reflective of a host of potential naturally arising fluctuations in activity external to this model microcircuit (***Kriener et al., 2008; Roelfsema et al., 2004; Doiron et al., 2016; Honey and Valiante, 2017; Lee et al., 1998; Stark et al., 2008***) and known to induce oscillatory synchrony (***Butler et al., 2017***). This correlation was found to significantly affect the SLE rate, as quantified in Figure 4**A**. With default values of each adaptation gain, all tested values of *c* > 0.1 yield significant increases in the SLE rate compared to the minimally correlated case of*c* = 0.01 (*p* < 0.05, two sample t-test). Furthermore, any increase in *c* of at least 0.3 also yielded a statistically significant increase to the SLE rate (*p* < 0.05, two sample t-test).

**Figure 4.**
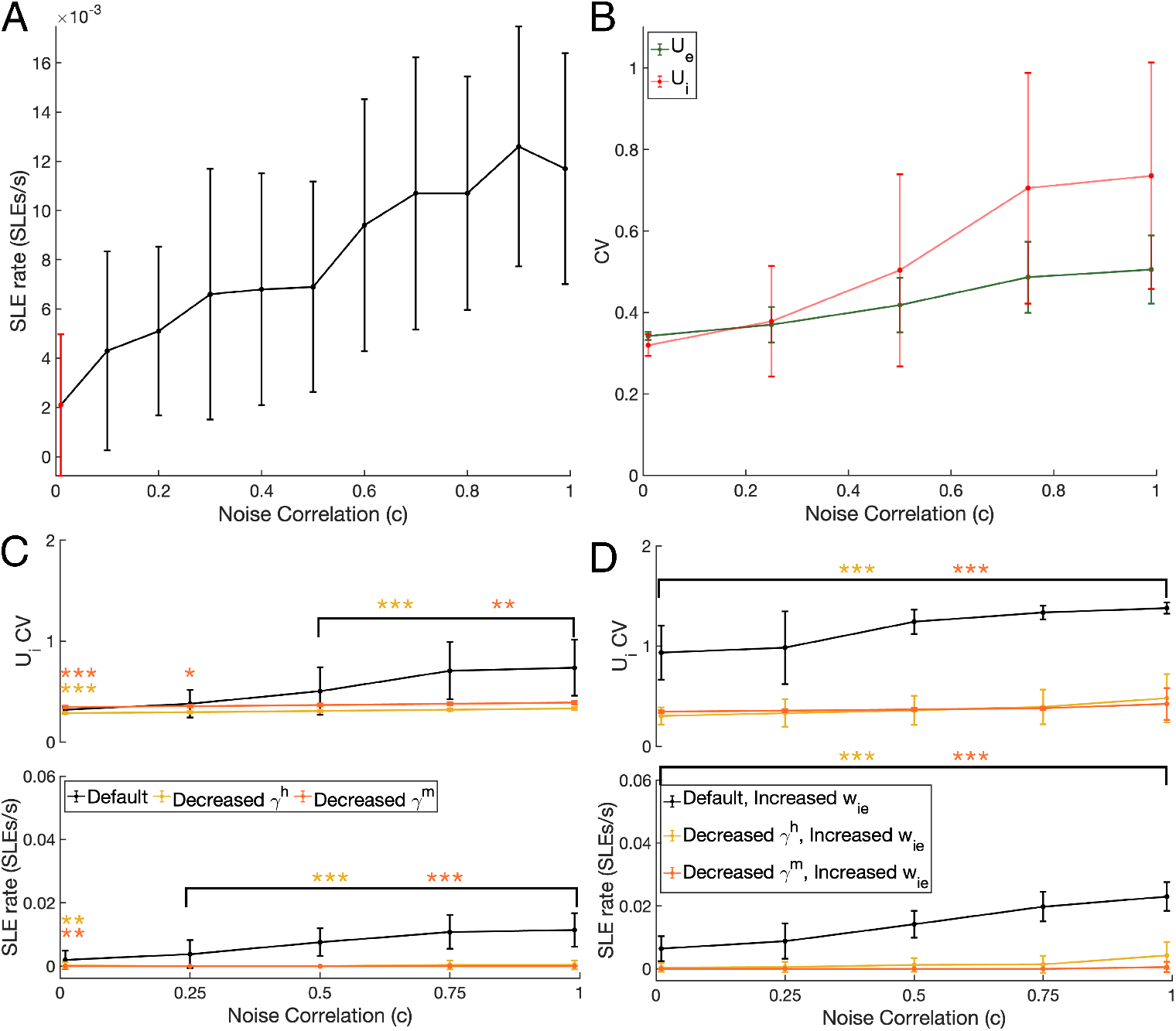
Model interrogation with varying degrees of input correlation identifies inhibitory population activity variability as a robust biomarker epileptogenic changes increasing SLE rate. **A**: SLE rate (SLEs/s) as a function of input correlation *c*. Plotted values are mean ± SD over 50 independent 200 s simulations. SLEs are significantly more frequent for each displayed *c* value in comparison to the red *c* = 0.01 case (*p* < 0.05, two sample t-test). **B**: Excitatory (*U*_*e*_, left) and inhibitory (*U*_*i*_, right) population variability (via the coefficient of variation, abbreviated CV) as a function of noise correlation *c* in the default model. Plots are mean ± one SD over 25 independent 200 s simulations, with the variance calculated excluding high-frequency firing as described in the Materials and Methods. Variance increases in both populations as a function of *c*, with more pronounced effects in the inhibitory population. **C**: *U*_*i*_ variability (top) and SLE rate (bottom) in the default model compared to two manipulations with minimal seizure-like activity, one with γ^*h*^ = 0.9 and the other with γ^*m*^ = 40.0. Plots are mean ± one SD over 25 independent 200 s simulations. Both alterations exhibit significantly reduced variability for *c* ≥ .5 and minimal response to increasing *c*, with these changes clearly correlated with changes in the SLE rate. Statistically significant differences, as compared to the default model, calculated using the two-sample t-test and denoted by conventional * symbols. **D**: The ictogenic effect of increased *w*_*ie*_ in the default model is significantly diminished with decreases to γ^*h*^ and γ^*m*^ as in Panel **C**. Plots and significance as described for Panel **C**.

We hypothesized that these ictogenic consequences might be driven by pathological variability in the excitatory and/or inhibitory population response to correlated input. Indeed, an increased likelihood of high-amplitude activity caused by correlated input could trigger a state transition towards SLEs. We thus quantified the variability of the mean excitatory and inhibitory activity (*U*_*e*_ and *U*_*i*_, see Materials and Methods for details) as a function of input correlation exclusively during non-SLEs epochs to avoid potential confounds. To do so, we only considered the system’s baseline activity by excluding periods with high-frequency spiking activity indicating an SLE and/or inter-ictal like event (see Materials and Methods for details). This analysis, performed for our default parameter values, is presented in Figure 4**B**: we observe that the coefficient of variation (CV) of the inhibitory membrane potential increases more notably with *c* compared to excitatory cells. These results indicate that input correlation predominantly affects inhibitory activity, leading us to hypothesize that fluctuations in inhibitory activity might serve as a biomarker for SLE susceptibility.

As an initial test of this hypothesis, we asked how the relationship between input correlation and inhibitory population variability would change with varying h- and m-adaptation gains. Based on the analyses presented in Figure 3**A**, we tested two such cases reflecting reductions to γ^*h*^ and γ^*m*^ (the lack of subscript reflecting a uniform change in both excitatory and inhibitory populations) that yielded a near-zero SLE rate. We found that changes in variability in inhibitory cell activity mirrored SLE susceptibility: decreases in h- and m-adaptation suppresses the variability of inhibitory cell activity induced by correlated inputs. As illustrated in Figure 4**C**, the *U*_*i*_ CV in these anti-ictogenic scenarios was lower than in the default network for all tested values of *c* > 0.01.

We further confirmed the robustness of this measure as a biomarker for SLE susceptibility by subjecting the system to changes not to the h- and m-adaptation, but to the strengths of the connections in the E-I network topology. We independently altered each of the four synaptic weights (*w*_*ee*_, *w*_*ei*_, *w*_*ii*_, and *w*_*ie*_) to .5 and 1.5 times their default values and quantified the variance in *U*_*i*_. For *w*_*ee*_, *w*_*ei*_, and *w*_*ii*_ (Figure 5**A-C**) the results are intuitive: pro-ictogenic changes increasing network excitability (increasing excitatory synaptic strenghts or decreasing inhibitory synaptic strengths) yield increased *U*_*i*_ variability, while the reverse changes decrease *U*_*i*_ variability and the system’s sensitivity to changing *c*. In this light the results for *w*_*ie*_ (Figure 5**D**) are perhaps counter-intuitive, as increased inhibitory signaling leads to increased variance and SLE rate, while decreased inhibitory signaling leads to decreased variance and SLE rate as well as a reduced sensitivity to *c*. However, in the context of the SLE initiation mechanism described from analysis of Figure 1**D**, this in fact highlights a causal role for inhibitory signaling in generating SLEs in this system: stronger inhibitory-to-excitatory signaling will further hyperpolarize the excitatory population following a period of inhibitory hyperactivity, exacerbating the post-inhibitory hyperexcitability of the excitatory population necessary to trigger SLEs.

**Figure 5.**
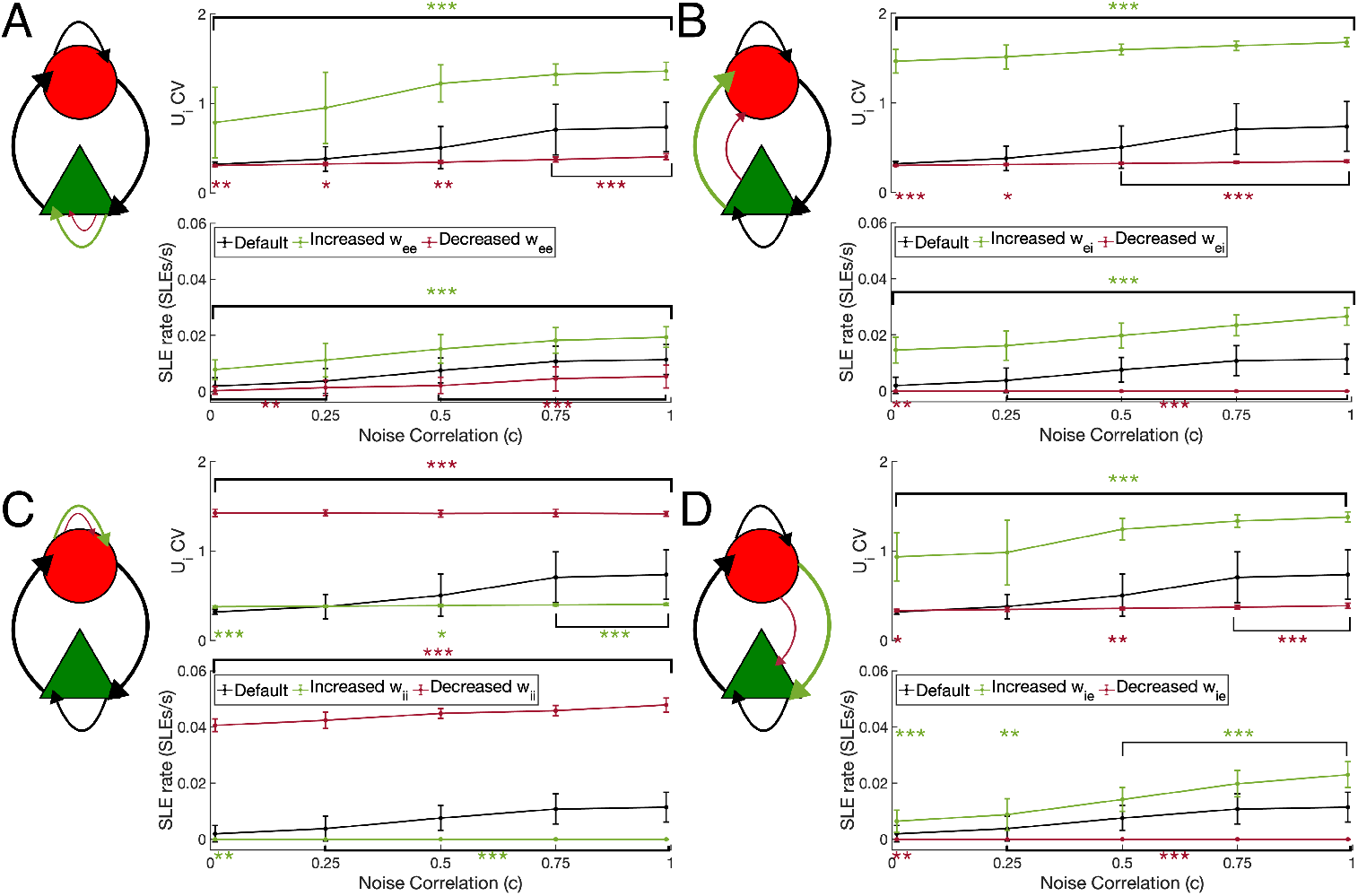
Alterations to network topology yield correlated changes to *U*_*i*_ CV and SLE rate reinforcing the viability of the *U*_*i*_ CV as a biomarker for SLE susceptibility. **A-D**: Analysis of *U*_*i*_ variability for increases (1.5 times the default value) and decreases (0.5 times the default value) in the synaptic weights. Plots as in Figure 4**C**. Changes in *w*_*ee*_ **(A)**, *w*_*ei*_ **(B)**, and *w*_*ii*_ **(C)** are intuitive: increasing the amount of excitation or decreasing the amount of inhibition in the system will increase population variability amongst the inhibitory neurons, while changes decreasing this variability also reduce the response to increasing *c*. Changes in *w*_*ie*_ **(D)** oppose this trend: increasing the strength of inhibition significantly increases the variability and its dependence upon *c*, illustrating both the vital role played by inhibitory signaling in the modeled seizure-like activity and also the reality that the modeled system is in a pathological state. Statistically significant differences, as compared to the default model, calculated using the two-sample t-test and denoted by conventional * symbols.

Collectively, these analyses provide strong support for the *U*_*i*_ CV as a reliable biomarker for seizure susceptibility in this model. Notably, this biomarker has a biophysical foundation as a measure of the system’s capacity to decorrelate inputs: systems vulnerable to SLEs are more prone to high amplitude fluctuations in inhibitory activity in response to highly correlated input, while systems resilient to SLEs will otherwise quench such fluctuations in the processing of said input (***Middleton et al., 2012***). This explanation is supported by an additional finding, that the effects of changing *w*_*ie*_ are notably diminished in the two anti-ictogenic scenarios studied in Figure 4**C** (Figure 4D). This serves to reconcile these results with the understanding that inhibitory-to-excitatory signaling decorrelates physiological network activity (***Renart et al., 2010; Middleton et al., 2012***): this model is specifically of a pathological system, mirroring pro-ictogenic overexpression of h- and m-channels, unable to respond to input correlations in a healthy manner.

We finally asked whether this biomarker, and in turn the impairment of the processing of correlated input that it identifies, provides a viable mechanistic explanation for the ictogenic effects of varying adaptation gains observed in Figure 3. Setting the input correlation to *c* = .99 as in Figure 3 to make the system maximally susceptible to SLEs, we systematically varied the gains of h- and m-adaptation and examined the impact on the variability of inhibitory cell activity. Increased 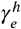 was found to monotonically increase the variability in *U*_*i*_ (Figure 6**C**), mirroring changes seen in Figure 3**D** where 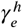 increases the SLE rate. Similarly, we observe notable changes in *U*_*i*_ variability as a function of 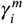 around the model’s default value (highlighted in red): decreased 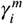 yields notable decreases in the *U*_*i*_ CV compared to the default scenario, while increased 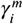 yields a notable increase in the *U*_*i*_ CV compared to this default (Figure 6**B**). This reflects the fact that a minor decrease in 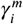 from its default value eliminates nearly all SLEs (Figure 3**D**), while a small increase leads to a pronounced increase in the SLE rate. Notably, these results showcase that the ictogenic consequences of h-and m-channel overexpression are not achieved directly through an increase in single cell excitability or the strength of the external input; rather, these channelopathies interfere with the processing of highly correlated input (without a necessary change in this input’s amplitude), yielding a cascade of effects throughout the network that provoke an SLE

**Figure 6.**
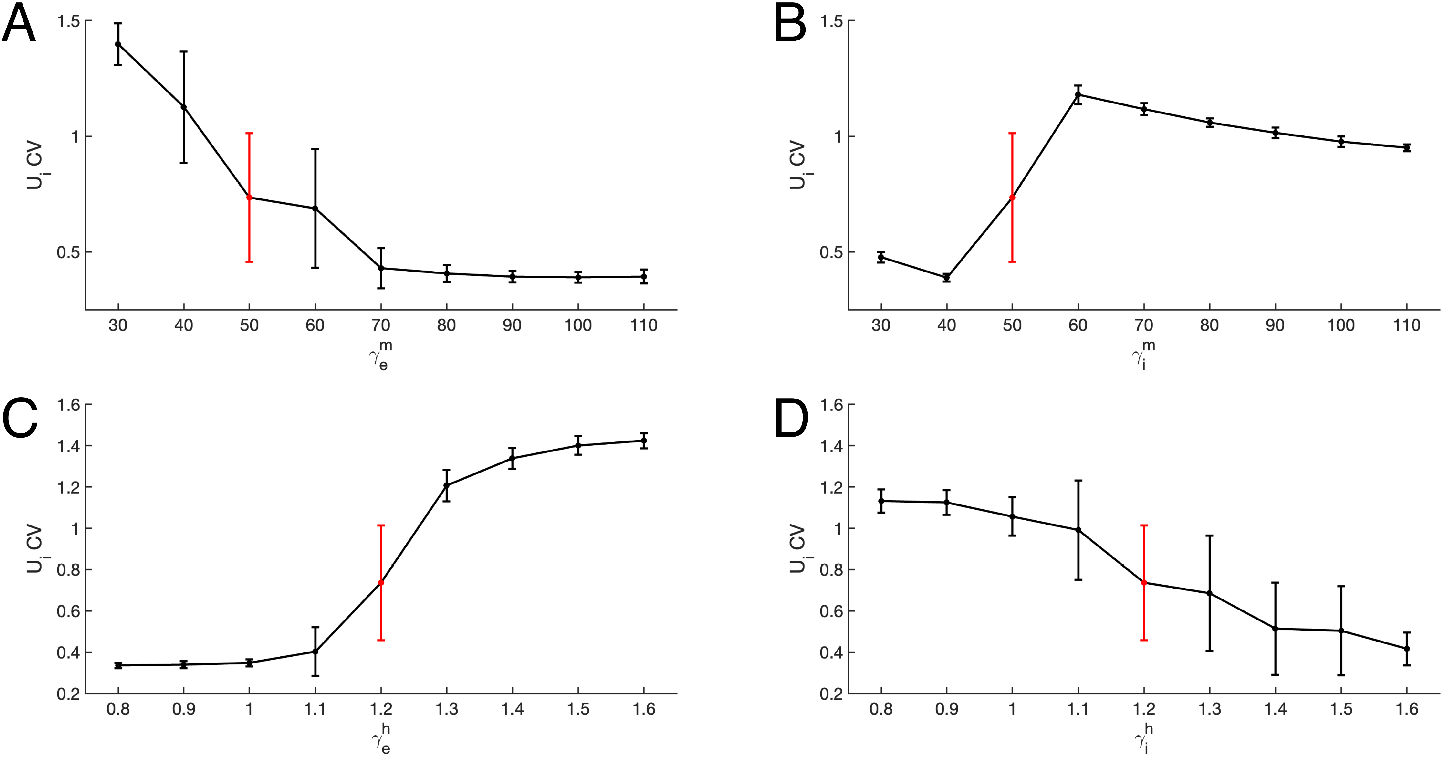
Ictogenic consequences of h- and m-channel overexpression can be mechanistically explained by the amplification of inhibitory variability triggered by a highly correlated input. **A-D**: *U*_*i*_ variability as a function of changing 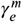 (Panel **A**), 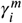 (Panel **B**), 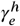 (Panel **C**), and 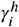 (Panel **D**). Plots are mean ± one standard deviation over 25 independent 200 s simulations with *c* = .99, with results from the default model parameters highlighted in red. Of note, intuitive hypothesized epileptogenic changes (e.g., increased 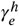 decreased 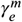 small increases in 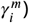 notably increase variability in *U*_*i*_.

More complex effects identified in Figure 3 can also be explained by this biomarker. Increases in 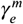 monotonically decrease the variability in inhibitory activity (Figure 6**A**), reflecting the pattern with default 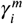 and increasing 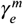 seen in Figure 3**C**. The more moderate changes of 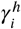 on the *U*_*i*_

CV (Figure 6**D**), not only conforms with the intuition from our analysis of Figures 1 and 2, but also the more moderate effect of γ^*h*^ on SLE rate in Figure 3**C**. We also note that the default model parameters’ uniquely high variance in the *U*_*i*_ CV and position at an inflection point in the the relationship between each parameter and the *U*_*i*_ CV serves as additional *a posteriori* justification for these choices, as this positioning promotes the unpredictable nature of SLE onset in this *in silico* model matching the clinical reality (***Kuhlmann et al., 2018; Mormann et al., 2007; Breton et al., 2020; Shayegh et al., 2014; Suffczynski et al., 2006; Snyder et al., 2008; Bauer et al., 2017; Colic et al., 2013***).

## Discussion

Through the presentation and analysis of a novel *in silico* seizure model whose epileptiform activity is dependent upon neuronal voltage homeostasis and spike-frequency adaptation, this work yields new insights into the epileptogenic effects of channelopathies affecting the h- and m-channels. Specifically, we describe a computationally-supported mechanism explaining the pro-ictogenic consequences of overexpression of the h- and m-channels (***Poolos, 2004; DiFrancesco et al., 2019; Noam et al., 2011; Bender et al., 2003; Dyhrfjeld-Johnsen et al., 2009; Niday and Tzingounis, 2018***), understudied channelopathies relative to the underexpression of these channels (***Poolos, 2004; DiFrancesco et al., 2019; Arnold et al., 2019; Dyhrfjeld-Johnsen et al., 2009; Lerche et al., 2001; Rogawski, 2000; Steinlein and Noebels, 2000; Byers et al., 2021***) that have effects more intuitively understood via changes to the E-I balance. We find that excessive h-adaptation promotes post-inhibitory hyperexcitability in excitatory neurons, which cannot be compensated for by inhibitory cells exhibiting excessive m-adaptation (Figure 1**D**). This impedes the ability of a neuronal micro-circuit to process a correlated input, resulting in more common high amplitude fluctuations in inhibitory activity that themselves can trigger SLEs. Critically, this mechanism indicates the ictogenic consequences of these channelopathies arises through a corruption of input processing, therefore addressing the paradoxical reality that both over-and under-expression of these channels can be ictogenic via two distinct mechanisms promoting seizure onset.

Notably, this mechanism parallels experimental findings that activation of parvalbumin positive (PV) interneurons at the seizure focus is seizure and synchrony-promoting via postinhibitory rebound (***Sessolo et al., 2015***). In this context, this work presents the homeostatic effects of the h-current (***Gasselin et al., 2015***) as a viable impetus for this postinhibitory rebound and subsequent ictogenesis, complementing the authors’ previous presentation of a “GABAergic initiation hypothesis” for seizure (***Rich et al., 2020a***). How this interacts with effects of the h-channel on input resistance (h-channel underexpression is thought to render neurons hyperexcitable by increasing the input resistance) is outside the realm of this study given our idealized neuron model; however, given our group’s previous focus on capturing the dynamics of the human h-channel (***Rich et al., 2021***), study of these interactions remains a ripe topic for future research. Similarly, this model is able to explain the less-intuitive ictogenic consequences of m-channel overexpression (***Niday and Tzingounis, 2018***) via its cell-type specific effects: excessive spike-frequency adaptation in inhibitory cells impairs their ability to restrain excessive excitatory activity, often outweighing the effects of such adaptation in the excitatory cells themselves. While underexpression of the m-current is intuitively related to ictogenesis via spike-frequency adaptation modulated effects on neuronal excitability and has motivated the upregulation of m-channels as a target for epilepsy treatment (***Shah et al., 2013; Seefeld et al., 2018; Kay et al., 2015; Rogawski, 2000***), our results indicate that such interventions should ideally be targeted to excitatory neurons.

The conclusions presented in this work are critically supported by this model’s capture of distinguishing nuances of seizure activity directly attributable to excessive h- and m-adaptation. Perhaps most notable amongst these features is the irregular nature of the SLEs, an analogue for the ongoing challenge of “seizure prediction” clinically (***Kuhlmann et al., 2018; Mormann et al., 2007***). Longer simulations of our model reveal that the non-periodic intervals between SLEs (an *in silico* analogue for inter-ictal intervals) are well-fit by a gamma distribution (Supplementary Figure S1), corresponding with experimental and clinical findings (***Breton et al., 2020; Shayegh et al., 2014; Suffczynski et al., 2006; Snyder et al., 2008; Bauer et al., 2017; Colic et al., 2013***). The model SLEs exhibit chirps (***Schevon et al., 2012; Cunningham et al., 2012***) and initial oscillatory frequencies in the 10-30 Hz range (***Cunningham et al., 2012; Li et al., 2008; Lee et al., 2000; Florez et al., 2013; Jiruska et al., 2013; Elahian et al., 2018; Velascol et al., 1999; Weiss et al., 2019; Perucca et al., 2014***), yielding a spectrogram in remarkable correspondence with experimental seizure recordings (***Florez et al., 2013; Huberfeld et al., 2015; Wendling et al., 2005; Schellenberg and Devergnas, 2020; Vuong et al., 2020***). Furthermore, examination of the spiking activity of individual neuronal units in the model, particularly during approximations of inter-ictal events, reveal that different excitatory neurons participate in each event and that inhibition plays a major role in their development (Supplementary Figure S2), matching experimental findings (***Feldt Muldoon et al., 2013; Muldoon et al., 2015***). Finally, the dynamics underlying the termination of our model SLEs mirrors “Fold Limit Cycle (FLC)” seizure offset as defined by ***Saggio et al***. (***2020***), the most common type of seizure offset identified in their recordings. Collectively, these findings indicate that this model not only is of critical use in the study of h- and m-channelopathies in epilepsy, but also in itself represents a crucial step forward in the fundamental endeavor of capturing the features differentiating seizure from other forms of oscillatory activity in *in silico* spiking neuronal microcircuits.

This assertion is further supported by the fact that the model realistically responds to a variety of epileptogenic insults beyond h- and m-channel overexpression, an important step towards computational models that reflect the diverse etiologies of epilepsy (***Jasper, 2012; Dehghani et al., 2016; Žiburkus et al., 2013; Rich et al., 2022; Sutula and Dudek, 2007; Cossart et al., 2001; Cobos et al., 2005; Arnold et al., 2019; Klaassen et al., 2006; Albertson et al., 2011; Rich et al., 2020a; Chang et al., 2018***). The model’s SLE rate can be affected solely by the correlation of a noisy input, reflecting a pathway to seizure driven by the inability to decorrelate a correlated input (***King et al., 2013; Sippy and Yuste, 2013***) and independent of changes to the amplitude of such input. However, the model still responds as expected to many changes to the E-I balance (***Dehghani et al., 2016; Žiburkus et al., 2013***). The ability to precisely and simultaneously study multiple, and potentially interacting, pro- and anti-ictogenic effects on distinct neuronal populations yields a range of novel insights into seizure onset speaking to this model’s potential impact on epilepsy research.

The unique role of inhibitory signaling in driving SLEs in this model (Figure 5**D**) is particularly salient given the lack of a clear consensus in the epilepsy literature as to the role of inhibitory neurons in seizure. While many studies have identified interneuronal hyperactivity prior to seizure onset and even implicated interneurons as serving a causal role in seizure initiation (***Sessolo et al., 2015; Muldoon et al., 2015; Elahian et al., 2018; Klaassen et al., 2006; Avoli and de Curtis, 2011; Avoli et al., 2016; Librizzi et al., 2017; Chang et al., 2018; Miri et al., 2018***), others contextualize inhibitory activity as primarily “restraining” seizure (***Schevon et al., 2012; Trevelyan and Schevon, 2013***) or focus on GABAergic signaling in the context of its tendency to become depolarizing during seizure propagation (***Chizhov et al., 2017, 2019; Ellender et al., 2014; Huberfeld et al., 2007***). Both these phenomena have strong experimental support, and compromised inhibition undoubtedly can be epileptogenic (***MacKenzie et al., 2016; Trevelyan and Schevon, 2013***); however, the apparent cap in the efficacy of pharmaceutical interventions directly affecting the E-I balance (***Brodie and Sills, 2011; Kwan et al., 2001***) implies that the role of inhibitory neurons in seizure is likely more nuanced and complex. Our model is strong computational evidence that SLEs with biophysically realistic features can arise with intact, completely inhibitory signaling that is at least complicit—and potentially causal—in SLE initiation.

Taken together, these conclusions reveal that h- and m-channel overexpression may corrupt the physiological role of inhibition in restraining seizure and processing correlated input. Indeed, activity in the cortex is typically uncorrelated (***Renart et al., 2010***) with inhibitory populations responsible for any necessary “decorrelation” (***Middleton et al., 2012; King et al., 2013; Sippy and Yuste, 2013***). However, the pathological nature of the default model studied here manifests in the amplification of the effects of input correlation (Figure 4) which is exacerbated with stronger inhibitory-to-excitatory signaling (Figure 5**D**). Collectively this implies that, in brain regions affected by epilepsy, seizure onset might not merely be driven by the failure of inhibition to decorrelate an upstream correlated input, but that the corruption of this processing plays a more causal role in seizure dynamics. This computationally-supported pathway merits rigorous interdisciplinary study and has the potential to recontextualize our understanding of the role of inhibition in seizure.

We note that the conclusions presented in this paper are conscientiously limited to seizure on-set rather than seizure propagation. This choice justifies the use of a small microcircuit comprised of 100 neurons and the omission of the effects of changes in the chloride reversal potential (***Lillis et al., 2012; Chizhov et al., 2019, 2017; Ellender et al., 2014***), which are most pronounced during seizure propagation (***Ellender et al., 2010; Chang et al., 2018***). Coupling multiple versions of this model together to study seizure propagation (***Proix et al., 2014; Jirsa et al., 2017***) and adding the associated changes to inhibitory signaling would be an interesting follow-up to this work. While such studies would ideally be performed with a reduced version of the model, the dynamical systems techniques used to do so in our previous work (***Rich et al., 2020b, 2022***) are not appropriate here given the sparse firing of the excitatory neurons outside of SLEs and the exaggerated stochasticity of the system’s dynamics. While the similarities (i.e., relative weighing of synaptic strengths) between this new model and those shown to exhibit multi-stability in our previous work (***Rich et al., 2020b, 2022***) are an indicator that multi-stability underlies the transition into SLEs studied here, confirmation of this structure will require new mean-field techniques.

Similarly, it is important to acknowledge that this model lies in an important middle-ground between primarily phenomenologically (***Jirsa et al., 2014; Kramer et al., 2005***) and biophysically (***Liou et al., 2020; Chizhov et al., 2022***) motivated models of seizure-like dynamics, and its applications must be correspondingly limited by the modeling focuses (***Almog and Korngreen, 2014***). While this microcircuit is comprised of individual spiking neurons and captures biologically-motivated stochasticity in the transitions into and out of SLEs, it also includes purposefully idealized models of the h- and m-currents. These models are designed only to capture key phenomenological effects of these currents of interest relative to the ictogenic effects of h- and m-channel overexpression. While future work will be required to determine whether the mechanisms proposed here are viable in more complex, biophysically-realistic models, the choices made in this study yield dynamics that are appropriately classified as “seizure-like” while maintaining sufficient mathematical and computational tractability, and thus are well justified given this study’s primary aims.

## Materials and Methods

### Model epileptogenic neuronal circuit

We present an epileptogenic spiking neuronal network model of recurrently connected excitatory and inhibitory neurons, inspired by numerous previous studies from our group (***Rich et al., 2020b, 2022***) and others (***Wilson and Cowan***, 1972; ***Kramer et al., 2005; Yang et al., 2005; Lytton, 2008; Cressman et al., 2009; Ullah et al., 2009; Kramer et al., 2012; Jirsa et al., 2014; Wendling et al., 2016; Buchin et al., 2016; Chizhov et al., 2018; Rich et al., 2020a; El Houssaini et al., 2020; Liou et al., 2020; Depannemaecker et al., 2021***). The model, which spontaneously transitions into SLEs, follows the general architecture of our previous work (***Rich et al., 2020b, 2022***) with the important addition of of spike-frequency adaptation and voltage homeostasis.

The spiking response of those neurons obeys the non-homogeneous Poisson process

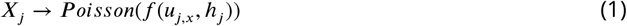

where 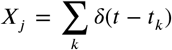 is a Poisson spike train with firing rate *f* (*u*_*j,x*_, *h*_*j*_) defined by the sigmoidal response function

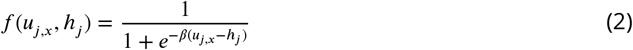

where *u*_*j,x*_ is the unitless membrane potential analogue for neuron *j*, where the index *x* = *e* when neuron *j* is an excitatory neuron while *x* = *i* when neuron *j* is inhibitory. In the equation above, *h*_*j*_ denotes the rheobase for neuron *j* that sets its excitability, while *β* refers to its response gain, chosen to be uniform across the network. Consequently, the probability of neuron *j* firing at time *t* is dependent upon *u*_*j,x*_ and is equal to

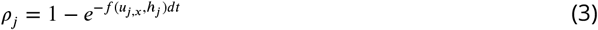

for *dt* sufficiently small. Heterogeneity in the network is introduced via the *h*_*j*_, which are chosen by independently sampling a normal distribution with standard deviation *σ*_*e,i*_ if the neuron is excitatory (*e*) or inhibitory (*i*). By default, *σ*_*e,i*_ = 0.01, modeling very low heterogeneity.

The dynamical system defining the temporal evolution of the membrane potential analogue *u*_*j,x*_ is given by the following:

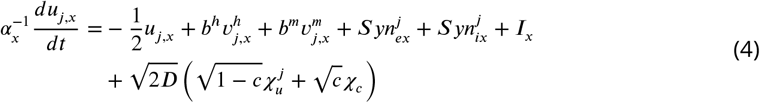

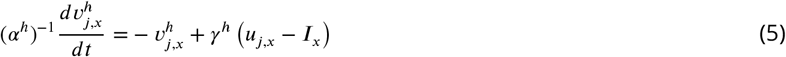

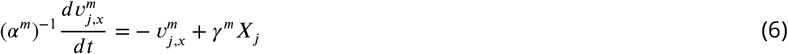

for *x* = *e* when neuron *j* is excitatory, and *x* = *i* when neuron *j* is inhibitory.

The variable 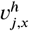 is our modeled voltage-homeostasis term, that evolves proportional to the difference between the current value of *u*_*j,x*_ and the bias current *I*_*x*_. Remembering that *b*^*h*^ < 0, this serves to excite the neuron when *u*_*j,x*_ < *I*_*x*_ for a prolonged period (given the slow time scale dictated by *α*^*h*^), and inhibit the neuron when *u*_*j,x*_ > *I*_*x*_ for a prolonged period. γ^*h*^ is the gain of this term, controlling the strength of this homeostatic drive. We refer to this term as “h-adaptation” considering that one well-studied effect of the h-current in neurons is voltage homeostasis (***Gasselin et al., 2015***).

The variable 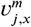 is our modeled spike-frequency adaptation term, which increases whenever neuron *j* spikes by a factor defined by the gain γ^*m*^ Remembering that *b*^*m*^ < 0, this serves to inhibit the neuron whenever there has been a spike in the recent past. We refer to this term as “m-adaptation” considering that a primary effect of the m-current is imbuing neurons with spike-frequency adaptation (***Benda and Herz, 2003***).

The terms 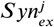 and 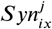 represent the synaptic inputs to the cell *j* from the excitatory and inhibitory neurons, respectively. For simplicity, we implement all-to-all connectivity. The stochastic nature of this system required a high number of simulation repetitions, so we limited our spiking network to 100 neurons, with the number of excitatory neurons *N*_*e*_ = 80 and the number of inhibitory neurons *N*_*i*_=20, matching the 4:1 ratio commonly studied in E-I networks (***Traub et al.,1997; Rich et al., 2017, 2018***). Accordingly, the post-synaptic inputs 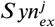 and 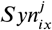 are given by

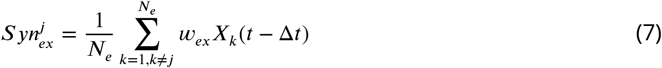

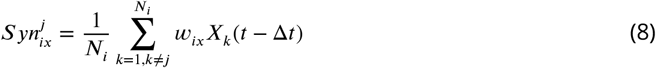

where *x* = *e, i* dependent upon whether neuron *j* is excitatory or inhibitory. Connectivity excludes self-synapses.

The term 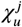represents an independent noise term sampled from *N*(0, 1) for each neuron, while*χ*_*c*_ *∈ N*(0, 1) is an additional independent noise term that is identical for all neurons. The value of *c* represents the noise correlation between neuronal units. When *c* = 0, the noisy input to each neuron is determined entirely by 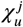 ; conversely, when *c* = 1, and the noisy input to each neuron isidentically determined by *χ*_*c*_.

For simplicity, we visualize the dynamics of the model via the population averages for the *u*_*x*_,*ν* _*h,x*_ and *ν* _*m,x*_ terms, simply calculated as 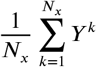where *x* = *e* for the excitatory population and *x* = *i* for the inhibitory population, and *Y* is a place holder for any of the 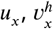, or 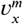 terms. These population averages are what are plotted in the visualizations of network dynamics presented in Figure 1, with the population averages of the *u*_*x*_ terms commonly referred to as *U*_*x*_. The *U*_*x*_ terms are also used for spectrogram analyses and analyses of the population variance (see below for details).

Parameters defining the system are found in Table 1. All random sampling is done in Python using numpy functions (***Harris et al., 2020***). Equations are integrated using the Euler-Maruyama method. In our simulations, Δ*t* = 0.1, scaled so that each time step Δ*t* represents 1 ms. The model is implemented in Python 3 and *will be made openly accessible via a lab GitHub repository at the time of final publication of this manuscript*.

**Table 1.**
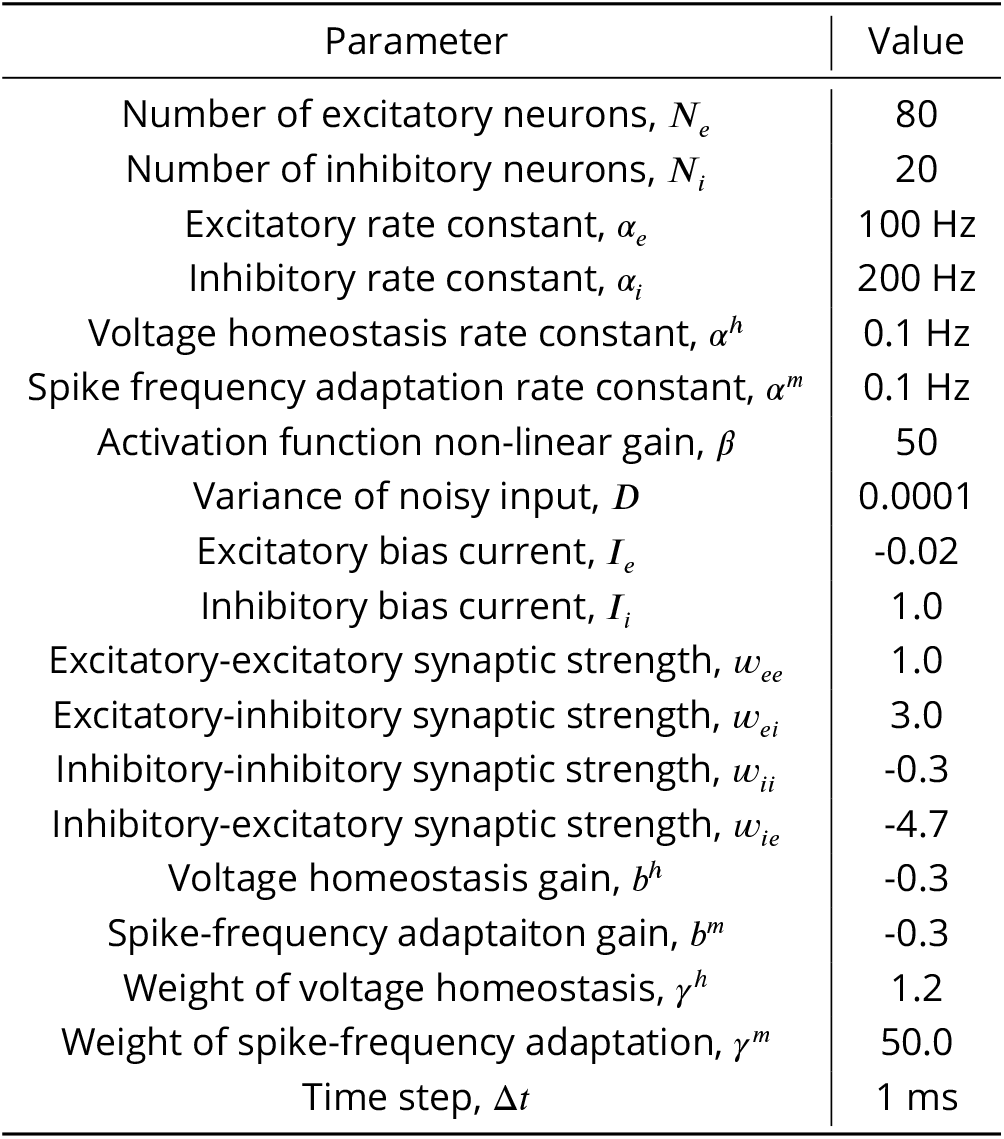
Default mean-field model parameters.

| Parameter | Value |
| --- | --- |
| Number of excitatory neurons, $N_e$ | 80 |
| Number of inhibitory neurons, $N_i$ | 20 |
| Excitatory rate constant, $\alpha_e$ | 100 Hz |
| Inhibitory rate constant, $\alpha_i$ | 200 Hz |
| Voltage homeostasis rate constant, $\alpha^h$ | 0.1 Hz |
| Spike frequency adaptation rate constant, $\alpha^m$ | 0.1 Hz |
| Activation function non-linear gain, $\beta$ | 50 |
| Variance of noisy input, $D$ | 0.0001 |
| Excitatory bias current, $I_e$ | -0.02 |
| Inhibitory bias current, $I_i$ | 1.0 |
| Excitatory-excitatory synaptic strength, $w_{ee}$ | 1.0 |
| Excitatory-inhibitory synaptic strength, $w_{ei}$ | 3.0 |
| Inhibitory-inhibitory synaptic strength, $w_{ii}$ | -0.3 |
| Inhibitory-excitatory synaptic strength, $w_{ie}$ | -4.7 |
| Voltage homeostasis gain, $b^h$ | -0.3 |
| Spike-frequency adaptation gain, $b^m$ | -0.3 |
| Weight of voltage homeostasis, $\gamma^h$ | 1.2 |
| Weight of spike-frequency adaptation, $\gamma^m$ | 50.0 |
| Time step, $\Delta t$ | 1 ms |

### Time-frequency analyses

Our primary tool for quantitatively assessing the dynamics of the model is time-frequency analysis using the *pspectrum* function in MATLAB (***MATLAB, 2019***). We perform this analysis on the mean activity of the excitatory population (*U*_*e*_), using the “spectrogram” mode with “TimeResolution” of 1.563 s and the “MinThreshold” option set to -17.5.

Thorough direct inspection of individual simulations revealed that SLEs were hallmarked by an increase in power in the 10-30 Hz band not seen during any other dynamics. This demarcates these events from brief instances of synchrony reminiscent of inter-ictal events and was consistent across our various manipulations to the model. We quantitatively identified such dynamics by: 1) Creating a time series of the summed power over this frequency range; 2) Smoothing this time series using MATLAB’s *smooth* function (***MATLAB, 2019***) with a “span” of 10; 3) Determining when this smoothed time series was > 10. Each continuous period for which this occurred is deemed an SLE and counted for the purposes of presenting the SLE rate.

### Additional *in silico* experiments and analyses

We present analyses of the coefficient of variance (CV) of *U*_*e*_ and *U*_*i*_ and how this changes as a function of *c* and other manipulations of the model. We calculate the CV only when the model is in a “baseline” state by excluding periods with any high-frequency spiking activity that could be indicative of an inter-ictal like event or an SLE. Given the effects that the various manipulations have on the baseline firing tendencies of the model, we accomplish this by: 1) Taking the spectrogram of the mean excitatory activity as usual; 2) Finding the maximum frequency for which the mean power over time is <0.001, which we’ll refer to as the “cutoff frequency”; 3) Creating a summed power time series from this cutoff frequency to 30 Hz (much like we did previously between 10-30 Hz for detecting SLEs); 4) Smoothing this power series as done in the detection of SLEs; 5) Excluding the time periods for which this smoothed power series is >0.01. We then calculate the CV only over the non-excluded times.

Additional data processing was required to derive the mean trajectory of the adaptation terms around SLEs presented in Figure 1**D**. After detecting SLEs as described above, we only considered SLEs beginning more than 50 s into the simulation to safely avoid any confounds by the the network’s initial state. We then extracted the mean activity (*U*_*e*_ and *U*_*i*_) and mean adaptation terms 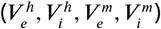 in three windows: the 5 s prior to detected SLE start, the duration of the SLE, and the 5 s after SLE end. The windows prior to and after SLE were directly averaged across the 57 SLEs. The time series during the SLE itself, given the variable duration of these events, were first normalized into time series of equal length (using the MATLAB (***MATLAB, 2019***) function *interp1* to interpolate these time series) and then averaged these processed time series which uniformly had 10000 time steps between the normalized “SLE Start” and “SLE End”. Means for each value outside of SLEs (the black dotted line with black shading representing ± SD in Figure 1**D**) was calculated by averaging the time-average of each time series between 50 s and simulation end (200 s) excluding SLEs. Average FRHs were created from 20 equally spaced intervals during each SLE (to account for the different duration of these events), as well as in the 5 s before and after each SLE.

### Significance testing

All tests of statistical significance were performed using the two-sample t-test via the *ttest2* function in MATLAB (***MATLAB, 2019***). A standard threshold of *p* < 0.05 is used to report statistically significant differences.

## Acknowledgments

We thank Dr. Frances Skinner for her insights and input on early versions of the model presented here.

## Supplementary Figures

**Supplementary Figure S1.**
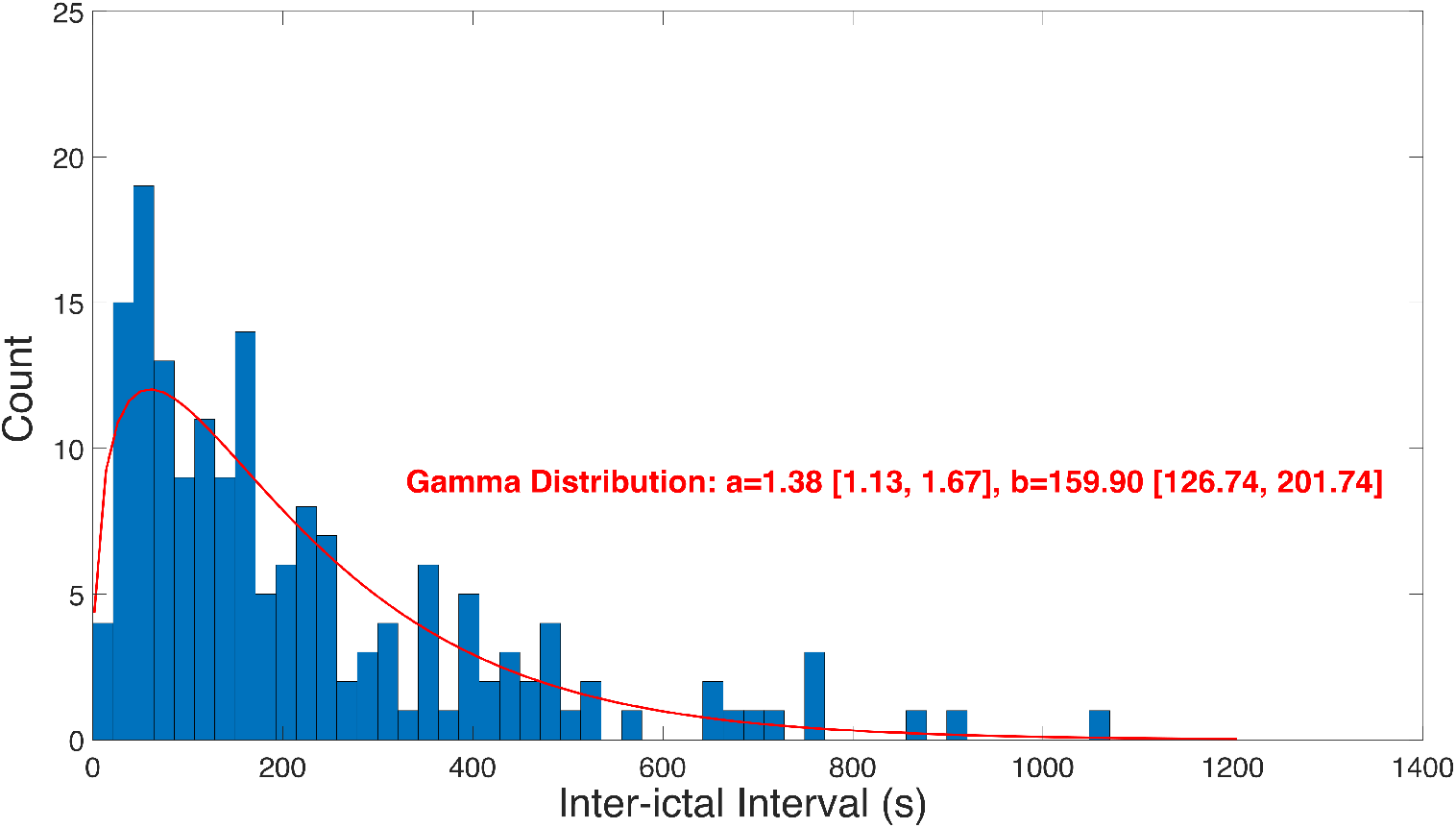
Intervals between SLEs are well fit by a gamma distribution. Histogram of intervals between SLEs taken from four 10,000 second simulations of the default model with *c* = .10 reveals the non-periodic nature of seizure onset. Data is well fit by a gamma distribution (red curve) with shape parameter *a* and rate parameter *b* (95% confidence interval in brackets).

**Supplementary Figure S2.**
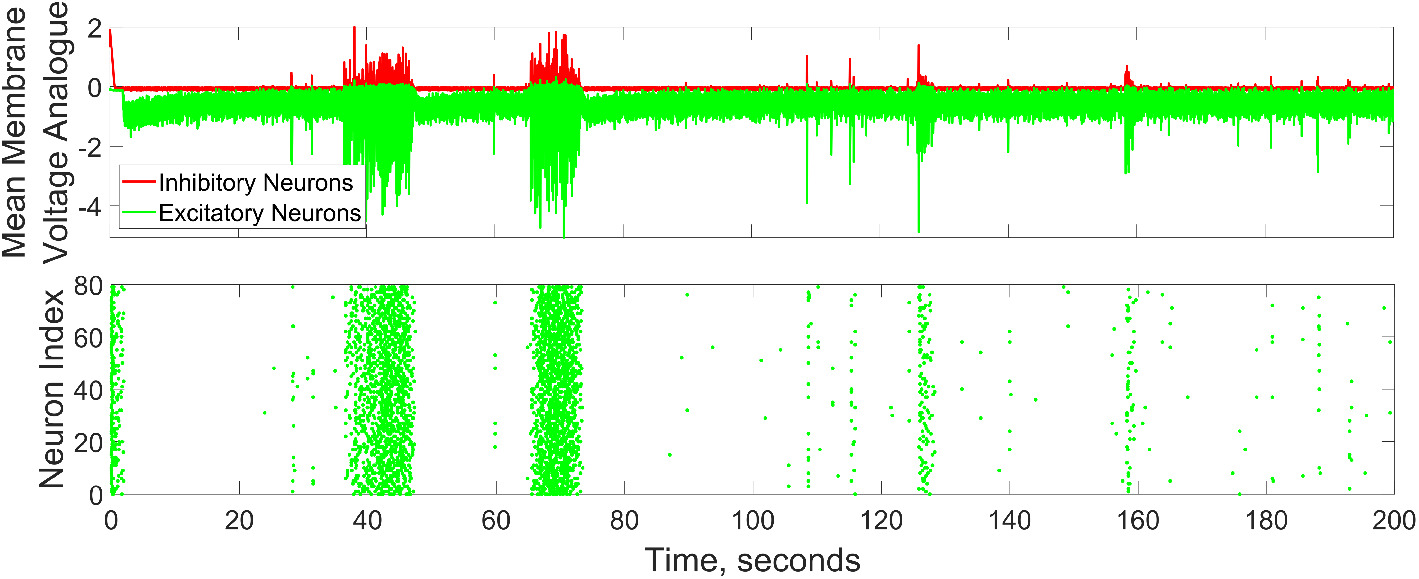
Example raster plot illustrating spiking activity of excitatory neurons. Raster plot corresponding to example simulation illustrated in Figure 1. Of particular note is the distinct populations of excitatory cells participating in “bursts” of activity reminiscent of inter-ictal spikes (between 100 and 160 s).

